# Segment-dependent phylogenetic resolution reveals geographic and host sampling gaps in Oropouche virus genomes

**DOI:** 10.64898/2026.11.05.764562

**Authors:** Rania Siam, Ali H. A. Elbehery

**Affiliations:** Ross University School of Medicine, Saint Michael, Barbados; Department of Microbiology and Immunology, Faculty of Pharmacy, University of Sadat City, Sadat City, Egypt

## Abstract

**Background:** Oropouche virus (OROV) has been reported in areas beyond its historical Amazonian range since 2023, accompanied by rapid growth in publicly available genomic data. However, the extent to which these data provide reliable phylogenetic and epidemiological resolution, and whether that resolution differs among genome segments, has not been systematically examined.

**Methods:** Nucleotide sequences annotated under *Orthobunyavirus oropoucheense* were retrieved from NCBI Virus (accessed 13 May 2025) and filtered to retain records at least 70% of the reference length for each segment, yielding 898 S, 831 M, and 832 L sequences. Segments were aligned independently with MAFFT, and maximum-likelihood phylogenies were inferred with IQ-TREE using ModelFinder and ultrafast bootstrap approximation. Trees were annotated with collection date, country, and host in iTOL.

**Results:** Phylogenetic resolution differed among segments. Contemporary Brazilian sequences formed a largely unresolved region in the S segment tree, whereas the M and L trees resolved recent clusters to differing degrees. Resolution did not track segment length: the M tree resolved contemporary structure more strongly than the longer L tree, and total tree length was approximately 3.5 times greater in the M tree despite near-identical taxon sampling. Sequences collected before 2010 were recovered in supported clusters in all three segment trees, although their configuration differed by segment. These sequences were also predominantly non-Brazilian and non-human, preventing collection date, geographic origin, and host from being separated as explanations for the observed clustering. The species-level query also retained three reassortant viruses, which may influence segment-specific topologies; sequences annotated to the United States were Iquitos virus rather than OROV. Two Italian sequences clustered with Cuban sequences across all three trees, concordant with published travel histories.

**Conclusions:** Publicly available sequences contain phylogenetic structure that differs among the S, M, and L segment trees, but interpretation is constrained by uneven temporal, geographic, and host sampling and by the inclusion of reassortant species in a species-level dataset. The concentration of sequences from Brazil and human hosts, together with limited representation of non-human hosts, restricts inference about OROV diversity and ecology. Broader and more balanced genomic surveillance would improve the ability to resolve OROV evolutionary and transmission patterns.

## Introduction

Oropouche virus (OROV) is an emerging arbovirus of the family *Peribunyaviridae*, genus *Orthobunyavirus* (species *Orthobunyavirus oropoucheense*), with a tri-segmented negative-sense RNA genome comprising small (S), medium (M), and large(L) segments. OROV circulates through a sylvatic cycle involving non-human primates, sloths, rodents, and possibly birds, and an urban epidemic cycle in which humans serve as the primary amplifying host. The definitive sylvatic vector has not been established, although *Aedes serratus, Psorophora cingulata*, and *Haemagogus tropicalis* have been implicated (Naveca et al., 2024). Urban transmission is attributed principally to the biting midge *Culicoides paraensis*, supported by repeated field isolation and epidemiological association during outbreaks (Pinheiro et al., 1981; World Health Organization, 2024) and by experimental vector competence data (Gallichotte et al., 2025). The contribution of mosquitoes is more limited than once assumed: across pre-2024 competence experiments, *Culicoides* spp. transmitted OROV at rates near 30%, whereas *Aedes* and *Culex* spp. were infected at rates below 20% and rarely transmitted OROV (Gallichotte et al., 2025).

OROV causes Oropouche fever, an acute febrile illness characterized by fever, headache, myalgia, nausea, vomiting, and, in some cases, neurological manifestations (Pinheiro et al., 1981; Wang et al., 2025). Because this presentation closely mimics other co-circulating arboviral diseases, diagnosis is frequently delayed or missed. Historically confined to Amazonian regions of South America, OROV has undergone significant geographic expansion since 2023, with outbreaks in Ecuador, Venezuela, Cuba, Panama, and several non-endemic Brazilian states (Bello-Rodríguez et al., 2025; Delatorre et al., 2025; Martins-Filho et al., 2025; Rodriguez-Morales et al., 2025). Risk mapping and ecological models indicate that deforestation, agricultural activity, and climatic shifts have contributed to this spread (Fischer et al., 2025), and phylogeographic reconstruction resolves a multi-scale process in which short-distance diffusive movement is punctuated by rapid long-distance jumps consistent with human-mediated dispersal (Tegally et al., 2026). International travel has introduced OROV to non-endemic regions, with imported cases reported in Italy, Portugal, and the United States (Morrison et al., 2024; Rossini et al., 2025; Zé-Zé et al., 2025).

The current epidemic is associated with a reassortant lineage that emerged in the Brazilian Amazon, combining S and L segments of Iquitos virus-like ancestry with a retained OROV-prototype-like M segment (da Silva Brito et al., 2025; Naveca et al., 2024). Genome segment reassortment has been a persistent driver of OROV evolution (Nunes et al., 2019; Tilston-Lunel et al., 2015), and genomic surveillance has documented autochthonous transmission in new locations including Rio de Janeiro and Pernambuco (Azevedo et al., 2024; Moreira et al., 2025; de Melo et al., 2026) as well as travel-associated international spread (de Melo Iani et al., 2025). Because the three genome segments can reassort, and because they differ substantially in length, segment-level analysis is necessary to characterize how contemporary strains relate to one another and to historical isolates.

Genomic surveillance underpins each of these insights, and its value depends on the composition of the sequence record available. This study examines the publicly available OROV sequence data through independent phylogenetic analysis of the S, M, and L segments, characterizing the relationships recovered from each segment, the structure separating contemporary from earlier isolates, and the geographic and host coverage of the record as it currently stands.

## Materials and Methods

### Sequence data retrieval and filtering

Nucleotide sequences of Oropouche virus (OROV) were retrieved from the NCBI Virus database by querying entries annotated under the species *Orthobunyavirus oropoucheense* (NCBI Virus, accessed 13 May 2025) (Supplementary Table 1). Note that any sequence classified under *Orthobunyavirus oropoucheense* was retained, including reassortant viruses. The frequency of each virus is included in Table 1. The dataset therefore represents a snapshot of the public sequence record on that date. Only sequences with clearly defined genomic segment assignment — small (S), medium (M), or large (L) — and annotated host information were retained.

**Table 1.** Species composition of the retrieved dataset, as annotated in GenBank. Iquitos, Madre de Dios and Perdões viruses are reassortants classified under *Orthobunyavirus oropoucheense* and were returned by the species-level query. All sequences were retained.

| **Species as annotated in GenBank** | **S segment** | **M segment** | **L segment** |
|---|---|---|---|
| *Oropouche virus* | 797 | 750 | 750 |
| *Orthobunyavirus oropoucheense (not further specified)* | 79 | 74 | 74 |
| *Iquitos virus* | 18 | 3 | 4 |
| *Madre de Dios virus* | 2 | 2 | 2 |
| *Perdões virus* | 2 | 2 | 2 |
| **Total** | **898** | **831** | **832** |

Reassortant species accounted for 22 of 898 S sequences (2.4%), 7 of 831 M (0.8%), and 8 of 832 L (1.0%). The remainder were annotated either as Oropouche virus or under the species name alone (Table 1). Because these reassortants are historical isolates and several derive from non-human hosts, their retention is relevant to the interpretation of the temporal and host patterns reported below, and is addressed there.

Length-based filtering was applied to exclude incomplete records. Sequences were retained if their length was at least 70% of the reference length for the corresponding segment. Reference lengths were determined from the most frequent (modal) length among entries flagged as complete in GenBank for each segment. The S segment reference length was 946 nt, with sequences ≥663 nt retained (n = 898); the M segment reference was 4,385 nt, retaining sequences ≥3,070 nt (n = 831); and the L segment reference was 6,852 nt, with sequences ≥4,797 nt retained (n = 832). The shortest retained sequences were 663, 3,824, and 5,551 nt for S, M, and L respectively, so the length filter was binding only for the S segment.

Country metadata coverage varied by segment, and the number of sequences available per country was not identical across the three datasets (Supplementary Table 1). All accessions, with segment, collection date, country, and host, are listed in Supplementary Table 1.

### Sequence alignment

Multiple sequence alignments were generated independently for each genomic segment using MAFFT v7.471 (Katoh & Standley, 2013). The --auto option was used, which selected FFT-NS-2 for all three datasets given their size. Alignments were inspected manually for quality and consistency. The resulting alignments comprised 966 columns for the S segment, 4,518 for M, and 6,858 for L.

### Phylogenetic analysis

Maximum-likelihood phylogenies were inferred for each segment using IQ-TREE v2.4.0 (Minh et al., 2020). Substitution models were selected by ModelFinder using the Bayesian Information Criterion. For all three segments the best-fit model identified by ModelFinder Plus was TIM3+F+I+R4. Branch support was assessed by ultrafast bootstrap approximation with 1,000 replicates; clusters with ultrafast bootstrap values ≥95% were considered well supported. Trees were inferred without an outgroup and were not midpoint rooted; therefore, they should be regarded as unrooted phylogenies. Total tree length and summed internal branch length were recorded from the IQ-TREE outputs for each segment. Branch lengths are expressed in substitutions per site and are therefore independent of alignment length, so total tree length is directly comparable between trees inferred from the same taxa under the same substitution model. Quantitative comparison is made between the M and L trees, which contain 831 and 832 sequences respectively; the S tree contains 898 sequences and is not compared on this basis. These summaries describe the total inferred divergence of each tree and are not evolutionary rate estimates, as no molecular clock was applied.

### Metadata extraction and tree visualization

Collection date, country of origin, and host were extracted from GenBank records and used to annotate the trees. Visualization and annotation were performed in the Interactive Tree of Life (iTOL) v7.1.1 (Letunic & Bork, 2024); the most recent published description of the software documents version 6.

Sequences were classified by collection date as pre-2010 (collected before 1 January 2010), contemporary (collected on or after 1 January 2010), or undated (collection year unavailable). In practice, the contemporary set is dominated by epidemic years: sequences from 2023 and 2024 account for approximately 97% of contemporary sequences in each segment.

### Use of generative AI

During the preparation of this manuscript the authors used Claude (Anthropic) to assist with literature searching, reference formatting, drafting and editing manuscript text, and checking the internal consistency of the written paper against the tables, supplementary metadata, and reported figures. The tool had no access to raw sequence data and performed no sequence retrieval, alignment, phylogenetic inference, or other analysis; all data generation and analysis were performed by the authors. After using this tool, the authors reviewed and edited the content as needed and take full responsibility for the content of the publication.

### Limitations of the dataset

The dataset is strongly skewed in both time and geographic origin, as quantified in the Results, with the majority of sequences collected during 2023–2024 and originating from Brazil. This distribution is likely influenced by sequencing effort and should not be interpreted as a proxy for viral abundance. Observed tree structure therefore reflects sampling intensity alongside underlying viral diversity. Because the public sequence record does not provide a basis for correcting these sampling imbalances, the analyses characterize the observed sequence record without attempting to adjust for them.

The dataset represents the public sequence record as of 13 May 2025. Deposition has continued since that date, including sequences from countries underrepresented here: 39 genomes from Loreto, Peru (GenBank PX608145–PX608183) and nine from Venezuela (PX494099–PX494101, PX495048–PX495050, PX495206–PX495208) were released subsequently. These additions are modest relative to the dataset analyzed, are concentrated in human-derived sequences, and would reduce but not remove the geographic imbalance described above; they would not alter the scarcity of sequences from non-human hosts and vectors. We therefore report the May 2025 snapshot while noting that the record continues to grow.

## Results and discussion

### Composition of the retrieved dataset

After length filtering, 898 S, 831 M, and 832 L segment sequences were retained. Collection year was available for 886 (98.7%) S, 825 (99.3%) M, and 826 (99.3%) L sequences. Country of origin was recorded for 850 of 898 S sequences (94.6%) and for all M (831) and L (832) sequences.

The dataset is heavily skewed in both time and space. Sequences collected in 2024 account for 493 (55.7%) of S, 538 (65.2%) of M, and 539 (65.3%) of L sequences with known collection dates, with a further 216 sequences from 2023 in each segment. The period from 1960 to 2009 contributes 155 S but only 49 M and 49 L sequences. The intervening years from 2010 to 2022 are represented by only 22 sequences in each segment, so the dataset is effectively bimodal in time, with a near-absent middle period separating the historical isolates from those of the current epidemic. Brazil accounts for 783 of 850 S sequences with known countries (92.1%), 744 of 831 M (89.5%), and 744 of 832 L (89.4%). Countries other than Brazil contributed 67 S-segment, 87 M-segment, and 88 L-segment sequences. Seven countries are represented by one or two sequences each in every segment.

This distribution is likely influenced by sequencing effort rather than reflecting viral abundance alone, and it conditions interpretation of all results below. In a dataset in which nine of ten sequences derive from one country and approximately two thirds from one year, observed tree structure is influenced strongly by where and when sampling occurred and cannot be interpreted independently of these sampling patterns. We report the observed patterns with this limitation stated rather than controlled, as the public sequence record offers no means of correcting it.

Country metadata were complete for the M and L datasets and available for 94.6% of S sequences. The number of sequences contributed per country nonetheless differs between segments: Colombia contributes 10 S sequences but 45 M and 45 L, whereas Peru contributes 38 S but 23 M and 23 L. Sequence counts per country are comparable between the M and L datasets, though segment records cannot always be traced to a common isolate.

The retrieved dataset is also not composed solely of OROV. The species-level query returned sequences of three reassortant viruses alongside OROV and records annotated only to the species name (Table 1), with reassortant species accounting for 22 of 898 S sequences (2.4%), 7 of 831 M (0.8%), and 8 of 832 L (1.0%). Species annotation also affects the interpretation of individual accessions: the sequences annotated to the United States in this dataset are Iquitos virus rather than OROV. Retrieval at species level therefore returns records that are not OROV sensu stricto and that cannot be identified as such without inspecting the species annotation of each accession. This is relevant to any analysis that treats a species-level query as equivalent to a query for OROV, and it is the reason the United States sequences are not interpreted as travel-associated OROV introductions below.

### Resolution of contemporary sequences is segment-dependent and does not track segment length

Phylogenetic resolution varied considerably between the three segment trees. In the S segment tree, the majority of contemporary sequences collapsed into a single unresolved fan occupying approximately one third of the topology. Branch lengths are ignored for effective visualization (Figure 1a). Within this region the tree provides insufficient resolution for reliable clustering inference; the resolved structure in the S tree is confined to the earlier and predominantly non-Brazilian isolates. This is consistent with the lower informational content of the shorter S segment, whose alignment comprises 966 columns, across which contemporary Brazilian sequences are near-identical. Differences in sequence variation, sampling composition, and data quality may also contribute to the segment-specific patterns observed.

**Figure 1.**
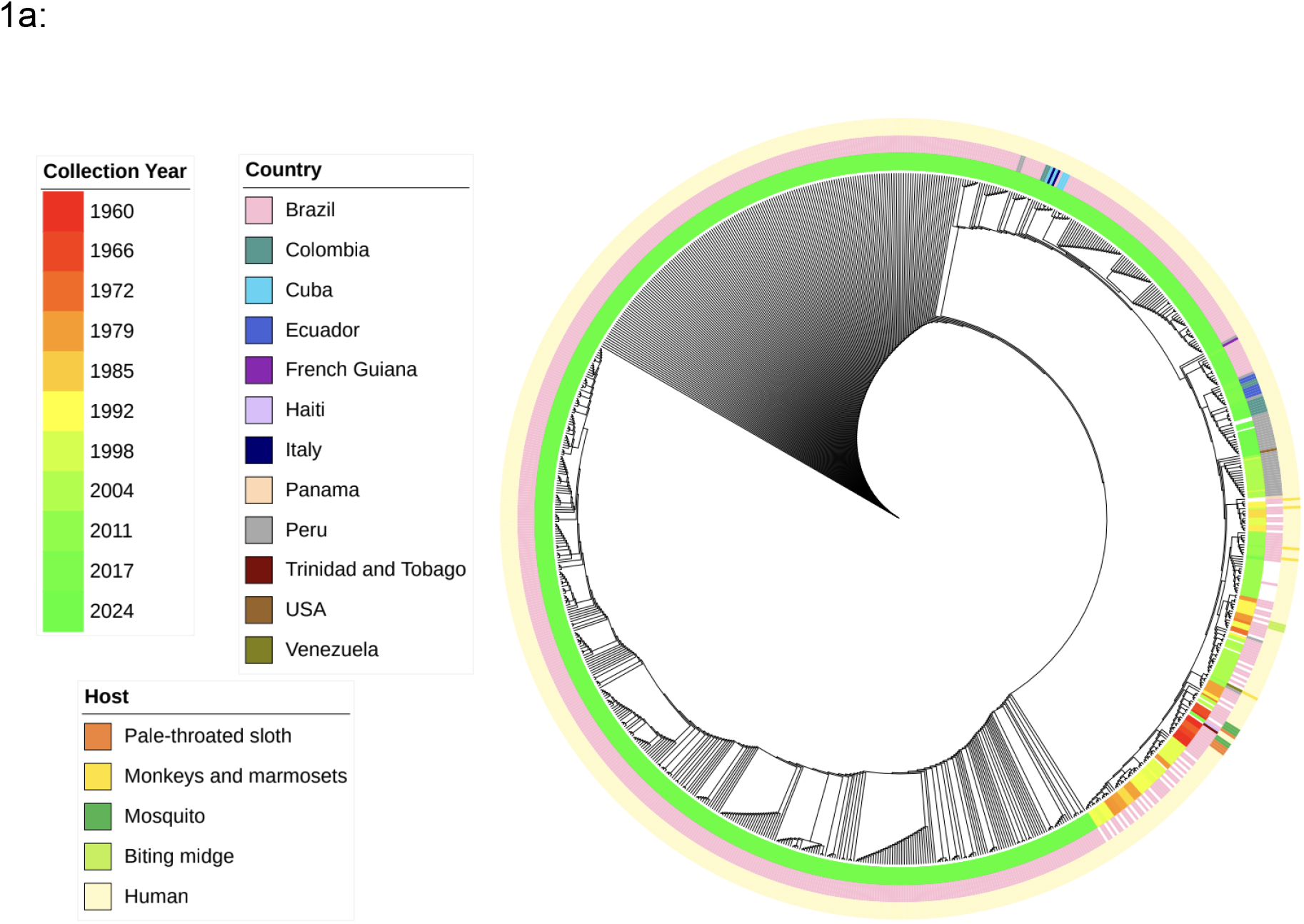
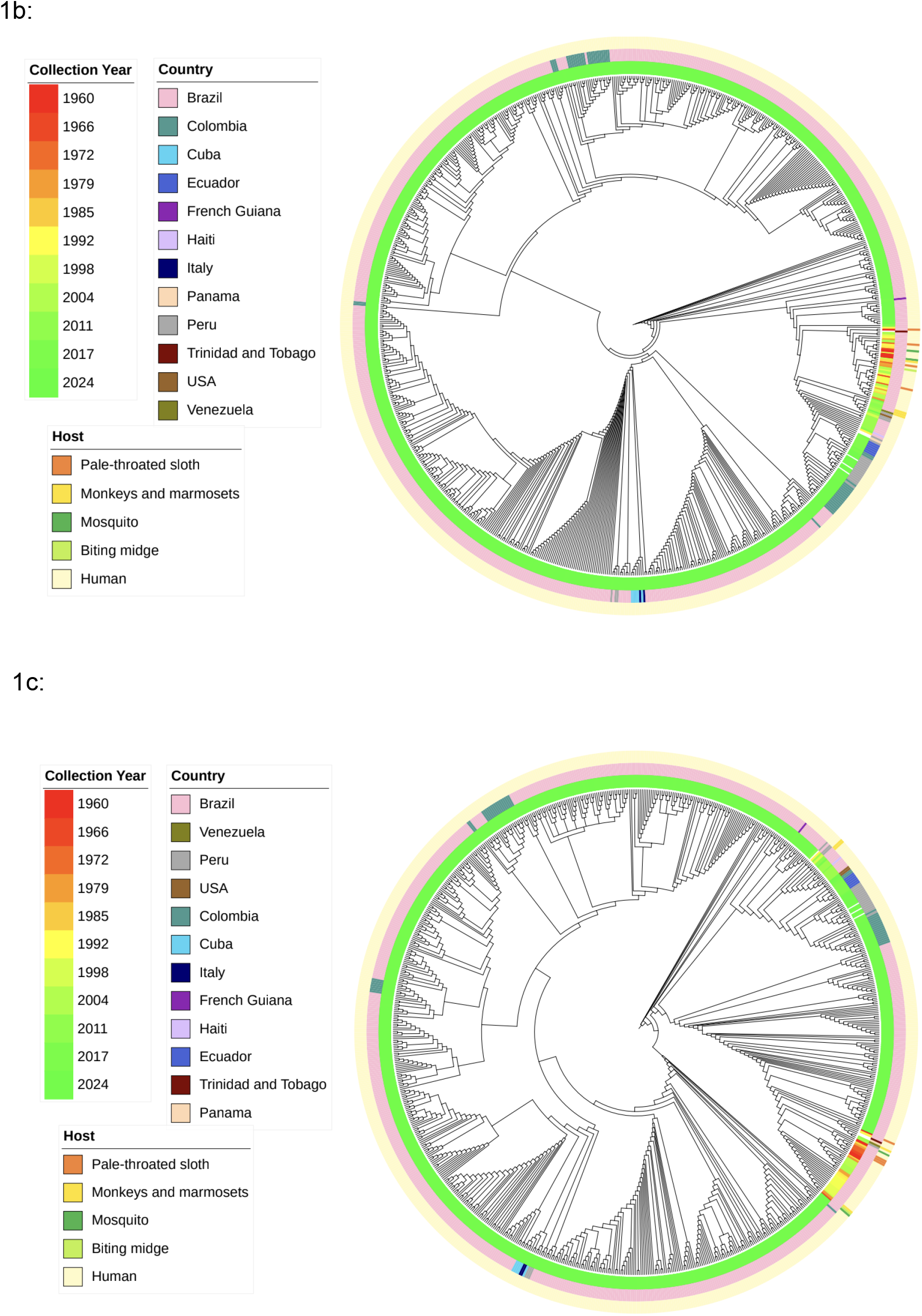
Maximum-likelihood phylogenies of the three Oropouche virus genome segments. (a) S segment (n = 898), (b) M segment (n = 831), (c) L segment (n = 832). Trees were inferred independently for each segment in IQ-TREE v2.4.0 under the TIM3+F+I+R4 substitution model selected by ModelFinder, with branch support assessed by ultrafast bootstrap approximation (1,000 replicates). Trees were inferred without an outgroup and are shown unrooted; the circular layout is used for display only and does not imply a root or a direction of evolution. Concentric annotation rings, from innermost to outermost, indicate collection year (colour gradient, 1960–2024), country of origin, and host. In (a), contemporary sequences collapse into a single largely unresolved region occupying approximately one third of the topology, with internal branch lengths approaching zero. Composition and branch support of the clusters referred to in the text are given in Table 2.

The M and L segment trees, whose alignments comprise 4,518 and 6,858 columns respectively, retain internal structure across contemporary sequences, resolving multiple distinct recent clusters rather than a single unresolved cluster (Figure 1b, c). Branch lengths in these regions are not depicted. In the M tree, a single cluster contained 685 of 776 contemporary sequences (88.3%) with 98% ultrafast bootstrap support. In the L tree, the corresponding node contained 518 of 777 contemporary sequences (66.7%) with 86% support; more inclusive clusters within the L tree were only weakly supported, with ultrafast bootstrap values of 24%, 11%, and 6% as cluster size increased. Contemporary cluster structure is therefore well resolved in the M tree, partially resolved in the L tree, and not resolved in the S tree.

Resolution does not, however, follow segment length. The L alignment is 52% longer than the M alignment and the two datasets contain comparable numbers of sequences (832 and 831), yet the M tree resolves contemporary structure more strongly than the L tree. Total tree length shows the same ordering (Table 2). Because branch lengths are expressed in substitutions per site, the total number is comparable between the M and L trees, which share essentially the same taxon sampling and were inferred under the same substitution model: the M tree totals 6.84 substitutions per site against 1.96 for L, approximately 3.5 times greater. Internal branches account for 92.0% of total tree length in the M tree and 58.4% in the L tree. The M segment therefore carries more inferred divergence, and more of it in internal branches, than the longer L segment.

**Table 2.** Branch-length summaries for the three segment trees. Branch lengths are in substitutions per site and are therefore independent of alignment length; total tree length is compared between the M and L trees, which have near-identical taxon sampling. Values are not evolutionary rate estimates.

| **Segment** | **Aligned sites** | **Total tree length** | **Internal branch lengths (% of total)** |
|---|---|---|---|
| S | 966 | 1.2018 | 0.5704 (47.5%) |
| M | 4,518 | 6.8381 | 6.2939 (92.0%) |
| L | 6,858 | 1.9588 | 1.1436 (58.4%) |

Several explanations may be compatible with the different topologies, but the present analysis does not distinguish among them. The M segment encodes the viral glycoproteins and the L segment the RNA-dependent RNA polymerase, so differences in functional constraints are one possible biological explanation. However, the dataset also retains reassortant species— Iquitos, Madre de Dios, and Perdões viruses—and their divergent segment histories may affect the inferred trees. Distinguishing these possibilities would require reanalysis excluding reassortant species, which was not performed. The supported conclusion is narrower: segment length alone does not explain the observed differences in phylogenetic resolution.

### Historical and contemporary sequences separate in all three segments, with segment-specific topology

Sequences collected before 2010 were recovered in supported clusters in all three segment trees (Figure 1a–c), although the form of that clustering differed between segments (Table 3). In the S tree, these sequences fell within a single cluster of 169 sequences (ultrafast bootstrap 91%), of which 155 were collected before 2010, five in 2010 or later, and nine were undated. In the M tree, the corresponding cluster of 99 sequences (98%) was mixed in composition, containing 49 pre-2010 and 44 contemporary sequences, but enclosed a nested subcluster of 38 sequences (100%) comprising 37 pre-2010 sequences and no contemporary sequences. In the L tree, pre-2010 sequences were distributed between two separate, non-overlapping clusters of 41 and 16 sequences, each with 100% support.

**Table 3.** Composition and branch support of the clusters containing pre-2010 sequences in each segment tree. Clusters are identified by their two terminal accessions. UFBoot, ultrafast bootstrap support (%).

| **Segment** | **Cluster (terminal accessions)** | **UFBoot** | **Total** | **Pre-2010** | **≥2010** | **Undated** |
|---|---|---|---|---|---|---|
| S | KJ866386.1 – HQ830446.1 | 91 | 169 | 155 | 5 | 9 |
| M | PQ801465.1 – PP477318.1 | 98 | 99 | 49 | 44 | 6 |
| M (nested subcluster) | PQ801465.1 – MG747597.1 | 100 | 38 | 37 | 0 | 1 |
| L (cluster 1) | MG747538.1 – KP052850.1 | 100 | 41 | 38 | 1 | 2 |
| L (cluster 2) | KP691615.1 – KC759125.1 | 100 | 16 | 11 | 4 | 1 |

Pre-2010 and contemporary sequences are therefore separated most cleanly in the M tree, where a fully supported subcluster contains no contemporary sequences, and least cleanly in the S tree, where support for the containing cluster (91%) falls below the threshold applied here and five contemporary sequences are included. The separation is consistent in direction across all three segments but is not recovered as a single, uniformly delimited clade.

These sequences differ from contemporary sequences in collection date, geographic origin, and host simultaneously, so the available data cannot disentangle which of these factors contributes to their separation. The temporal structure of the dataset contributes further. With only 22 sequences per segment collected between 2010 and 2022, the two groups are separated by a period that is sparsely sampled; some apparent discreteness may therefore reflect this gap rather than a discontinuity in the virus itself.

A further factor bears on the composition of these clusters. The species-level query returned sequences of three reassortant viruses classified under *Orthobunyavirus oropoucheense* (Table 1): Iquitos virus, Madre de Dios virus, and Perdões virus. All are historical isolates and several derive from non-human hosts. Sequences of these species are therefore expected to occupy the same region of each tree as the pre-2010 OROV sequences, and part of the separation described above may reflect divergence between virus reassortants rather than between time periods. The S segment is most affected, containing 18 Iquitos virus sequences against three in M and four in L; this asymmetry may also contribute to the differing cluster composition observed between segments.

Sequences from Peru, Ecuador, and Colombia are present in all three trees and are associated with the earlier, non-Brazilian cluster described above. These sequences do not, however, form a single clade. Colombian sequences in particular occupy at least three separate positions in the L tree and a comparable number in the M tree, suggesting that the sampled Colombian sequences occur across multiple lineages rather than representing a single introduction or a geographically coherent cluster.

Brazilian sequences formed multiple distinct clusters in the M and L trees. Sequences from other countries appear within Brazilian clusters at several positions. This is consistent either with cross-border transmission or with the resolution limits imposed by Brazil’s numerical dominance; the two cannot be distinguished with these data. Phylogenetic proximity is reported here as relatedness and is not interpreted as evidence of a transmission event.

### Gaps in non-human host representation

Non-human host sequences comprise 17 of 898 S sequences (1.9%), 11 of 831 M (1.3%), and 11 of 832 L (1.3%). At this level of representation, the data do not support inference of host-associated viral structure.

Almost all of these sequences fall within the pre-2010 clusters described above: 16 of 17 in the S tree, 11 of 11 in the M tree, and 11 of 11 in the L tree (nine in cluster 1 and two in cluster 2). They derive from sloths (*Bradypus tridactylus*), marmosets (*Callithrix penicillata* and *Callithrix* sp.), capuchin monkeys (*Cebus* sp.), mosquitoes (*Culex quinquefasciatus* and *Ochlerotatus serratus*, a synonym of *Aedes serratus*), and biting midges (*Culicoides paraensis*). The non-human-derived sequences in this dataset are therefore almost entirely historical, and their phylogenetic placement is inseparable from the collection dates and geographic origins of the same sequences. Detection of viral nucleic acid in an arthropod does not establish vector competence, and no inference regarding transmission capability is drawn.

The species annotation of these sequences warrants attention. Perdões virus was originally recovered from a marmoset and Madre de Dios virus from a capuchin monkey, and both are represented by two sequences per segment in this dataset (Table 1). A proportion of the non-human-derived sequences described here may therefore belong to reassortant species rather than to OROV itself, which would further limit what they can indicate about OROV host associations.

### A documented travel-associated introduction is recoverable from the public record

Two sequences from Italy clustered with Cuban sequences across all three segment trees (ultrafast bootstrap 99% in each), despite differences in phylogenetic resolution and country metadata availability among segments. Both sequences correspond to imported infections in travelers returning from Cuba, as independently documented in published case reports (Castilletti et al., 2024; Rossini et al., 2025). This is a specific instance in which phylogenetic placement is concordant with independently reported travel history; it does not establish transmission direction or validate travel-associated inference more generally.

Sequences annotated to the United States were not interpreted in this way. As described above, these accessions are Iquitos virus rather than OROV, so their phylogenetic placement reflects the identity of the virus sequenced rather than the travel history of an OROV infection. They are excluded from the observations reported here.

The Italian–Cuban clustering is a case-specific example of concordance between phylogenetic placement and published travel history. Because only two Italian sequences were available, it should not be taken as evidence that travel-associated introductions can generally be reconstructed from these data.

## Conclusion

This study examined the genomic diversification of Oropouche virus through independent maximum-likelihood phylogenies of the S, M, and L genome segments, using 898, 831, and 832 sequences, respectively, retrieved from the public record.

Three findings emerged. First, phylogenetic resolution differed remarkably among segments. The S segment showed limited resolution among contemporary Brazilian sequences, whereas the M and L trees resolved recent clusters to differing degrees. This ordering does not follow segment length: the M tree resolved contemporary structure more strongly than the longer L tree. Total tree length was approximately 3.5 times greater in the M tree than in the L tree despite near-identical taxon sampling, indicating that the M segment carries more inferred divergence than the longer L segment; whether this reflects functional constraint or the retained reassortant species was not resolved here. Second, sequences collected before 2010 were recovered in supported clusters in all three trees, though not as a single uniformly delimited clade: they formed one cluster in S, a fully supported subcluster nested within a mixed cluster in M, and two separate clusters in L. Because these sequences were also predominantly non-Brazilian and non-human, the effects of collection date, geographic origin, and host could not be disentangled; reassortant species retained from the species-level query may also contribute to this separation. Third, two Italian sequences clustered with Cuban sequences across all three segment trees, consistent with travel histories documented in published case reports. This is a case-specific concordance, not proof of transmission direction or a general measure of phylogenetic predictive performance. Sequences annotated to the United States were identified as Iquitos virus and were not interpreted as OROV introductions.

These results should be interpreted in the context of the available genomic data. The dataset is heavily weighted toward Brazil and toward sequences collected in 2023– 2024, reflecting sequencing effort rather than viral distribution; observed tree structure is influenced by this imbalance alongside underlying viral diversity. Sequences from non-human hosts number fewer than twenty per segment and fall almost entirely within pre-2010 clusters, so they do not support inference about host-associated viral structure. The analysis does not apply a molecular clock, test for selection, identify mutations associated with phenotype, or estimate evolutionary rates. The inclusion of reassortant species in the species-level dataset is an additional limitation because their segment histories may affect the observed topologies.

Within these limits, the analysis describes the phylogenetic structure recoverable from the public OROV-related sequence record as of 13 May 2025 and highlights its uneven geographic, temporal, and host coverage. The Italian–Cuban clustering provides one example of agreement between phylogenetic placement and independently reported travel history. However, sparse sampling outside Brazil, limited representation of non-human hosts, the near-absence of sequences from 2010–2022, and retention of reassortant species constrain inference about OROV diversity, ecology, and transmission. More geographically and temporally balanced sequencing, with explicit separation of OROV from related reassortant species, would strengthen future analyses.

## Declarations

### Data availability

All sequences analyzed are publicly available from NCBI Virus; accession numbers are listed in Supplementary Table 1.

### Author contributions

RS conceptualized the study, supervised the research, and wrote and edited the manuscript. AHAE conceptualized the study, performed the analyses, and edited the manuscript. Both authors read and approved the final manuscript.

### Ethics

This study analyzed publicly available sequence data only; no ethical approval was required.

